# *De novo* mass spectrometry peptide sequencing with a transformer model

**DOI:** 10.1101/2022.02.07.479481

**Authors:** Melih Yilmaz, William E. Fondrie, Wout Bittremieux, Sewoong Oh, William Stafford Noble

## Abstract

Tandem mass spectrometry is the only high-throughput method for analyzing the protein content of complex biological samples and is thus the primary technology driving the growth of the field of proteomics. A key outstanding challenge in this field involves identifying the sequence of amino acids—the peptide—responsible for generating each observed spectrum, without making use of prior knowledge in the form of a peptide sequence database. Although various machine learning methods have been developed to address this *de novo* sequencing problem, challenges that arise when modeling tandem mass spectra have led to complex models that combine multiple neural networks and post-processing steps. We propose a simple yet powerful method for *de novo* peptide sequencing, Casanovo, that uses a transformer framework to map directly from a sequence of observed peaks (a mass spectrum) to a sequence of amino acids (a peptide). Our experiments show that Casanovo achieves state-of-the-art performance on a benchmark dataset using a standard cross-species evaluation framework which involves testing with spectra with never-before-seen peptide labels. Casanovo not only achieves superior performance but does so at a fraction of the model complexity and inference time required by other methods.

## 1 Introduction

Tandem mass spectrometry provides a high-throughput framework for identifying and quantifying proteins in complex biological samples, but determining the exact protein content from observed mass spectra at scale remains a challenge. At the core of this challenge is the spectrum identification problem, in which we are given an observed mass spectrum and the associated mass and charge of the peptide (known as the *precursor*) that is responsible for generating the spectrum, and we must infer the amino acid sequence of the precursor peptide. The standard method for solving this problem is enumerative, scoring each observed spectrum with respect to a list of candidate peptides (i.e., peptides whose masses are close to the observed precursor mass associated with the spectrum) and reporting the best-scoring peptide-spectrum match (PSM) per spectrum.

However, the drawback to any database search methodology is that it requires that we specify *a priori* which peptides might occur in the sample. Such an approach is often sensible when analyzing samples from a species, such as human, with a well-characterized genome sequence. However, relying on a database prevents the detection of unexpected peptide sequences, such as those that arise from genetic variation. A sequence database also cannot be used for the analysis of some types of immunopeptidomics data [1], in antibody sequencing [2], or in vaccine development when searching for bacterial peptides present on the surface of infected cells [3]. Finally, constructing an accurate database for metaproteomic analyses, such as the human microbiome or environmental samples, is nearly impossible [4]. Such settings require *de novo* peptide sequencing from the acquired mass spectra.

Early *de novo* methods used heuristic search [5] or dynamic programming [6, 7, 8] to score peptide sequences against each observed spectrum. Machine learning has provided state-of-the-art performance on this task since 2015 [9], and recent methods employ deep neural networks [2, 10, 11, 12]. Although *de novo* search tools are improving, there is still a long way to go. The most recent report [10] suggests that state-of-the-art methods achieve peptide-level recall (i.e., the percentage of spectra with the correctly assigned peptide) of 39-60%, depending on the dataset. However, this percentage is calculated only with respect to ground truth spectra that, by definition, were previously identified with high confidence by a database search.

Additionally, all of these methods involve complicated modeling schemes (Table 1) featuring different neural networks for different sub-tasks such as convolutional neural networks (CNNs) for spectrum peak embedding and spectrum processing, and recurrent neural networks (RNNs) for peptide sequence processing. These methods also include complex post-processing steps that involve either matching the predicted peptide mass with the spectrum’s observed precursor mass using dynamic programming or refining low confidence predictions with a database search-like procedure. The necessity of discretizing the mass-to-charge (*m/z*) axis of the mass spectra also remains a setback for all existing methods except PointNovo, necessitating a trade-off between low binning resolution (hence low sequencing accuracy) and higher model complexity (hence longer inference time).

**Table 1:**
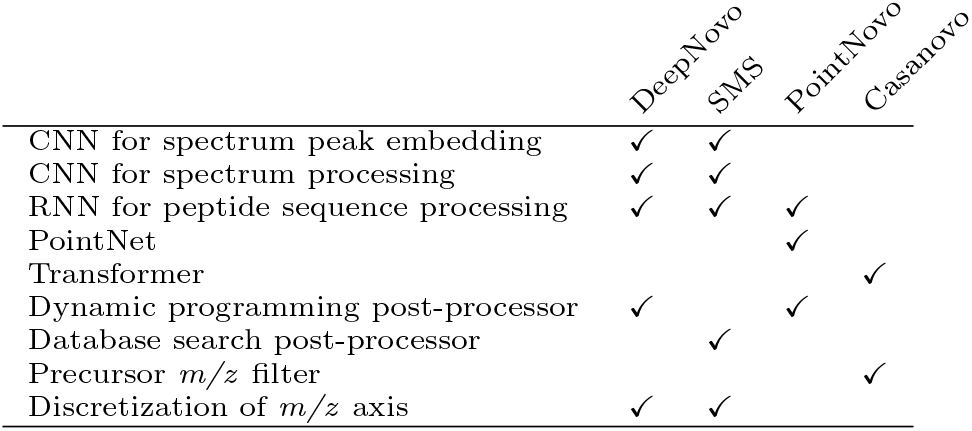
Comparison of deep learning methods for *de novo* peptide sequencing. Casanovo introduces a simpler yet more powerful architecture of a transformer for *de novo* peptide sequencing.

|  | DeepNovo | SMS | PointNovo | Casanovo |
| --- | --- | --- | --- | --- |
| CNN for spectrum peak embedding | ✓ | ✓ |  |  |
| CNN for spectrum processing | ✓ | ✓ |  |  |
| RNN for peptide sequence processing | ✓ | ✓ | ✓ |  |
| PointNet |  |  | ✓ |  |
| Transformer |  |  |  | ✓ |
| Dynamic programming post-processor | ✓ |  | ✓ |  |
| Database search post-processor |  | ✓ |  |  |
| Precursor $m/z$ filter | | | | ✓ |
| Discretization of $m/z$ axis | ✓ | ✓ | | |

In this work, we propose Casanovo, a transformer framework for *de novo* peptide sequencing (Figure 1). Casanovo uses the self-attention mechanism to translate directly from a variable-length sequence of observed spectrum peaks to a variable-length sequence of amino acids, analogous to a neural machine translation model in the natural language processing setting. Importantly, Casanovo takes individual spectrum peaks, together with the precursor mass and charge, as input, without resorting to discretization of the *m/z* axis, and learns to predict the generating peptide sequence in a supervised setting in which ground truth sequences are obtained with database search. Unlike existing methods, Casanovo does not employ an additional RNN to process peptide sequences and replaces the dynamic programming post-processing step with a simple delta mass filter, offering a simpler yet more powerful framework.

**Figure 1:**
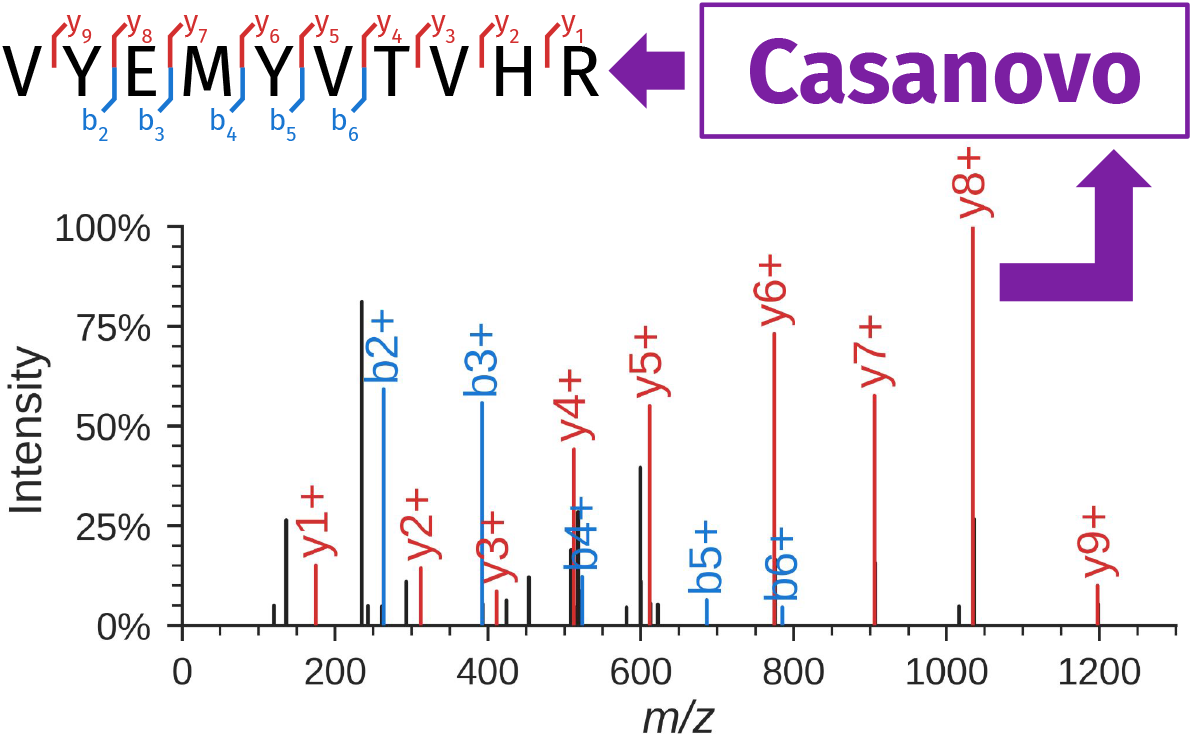
Casanovo performs *de novo* peptide sequencing. Casanovo takes as input an observed spectrum and produces the sequence of the generating peptide (e.g., VYEMYVTVHR). In the spectrum, peaks corresponding to b- and y-ions of the associated peptide are in color, and black peaks correspond to unexpected fragmentation events or noise. The spectrum annotation was created using spectrum utils [13].

We train and evaluate our model on a multi-species benchmark dataset using an established cross-validation framework which involves testing on spectra with never-before-seen peptide labels, through cross-species prediction. Our experiments show that Casanovo predicts peptide sequences with markedly higher precision relative to the state-of-the-art methods, DeepNovo and PointNovo, and does so using a model with fewer parameters and requiring much shorter inference time. Finally, we benchmark several variants of the model to demonstrate the robustness of some of our modeling choices.

## 2 Background: *De novo* peptide sequencing

Because tandem mass spectrometry analysis, and particularly *de novo* peptide sequencing, is rarely discussed in the machine learning literature, we begin with a brief overview of the problem domain. Additional background is available in several review articles [14, 15].

A tandem mass spectrometer measures mass-to-charge (*m/z*) ratios of charged peptides in a two-scan process. The first scan (MS1) measures the *m/z* of the intact peptide (also known as the precursor); the peptide is then fragmented and the resulting fragments are analyzed in a secondary scan (MS2). This MS2 scan is carried out on a population of (ideally) homogeneous peptide sequences, each of which is randomly fragmented at one location along the peptide backbone. As a result, the fragmentation scan contains peaks that correspond to prefixes (called *b-ions*) and suffixes (*y-ions*) of the peptide, each with an associated charge state. Thus, the primary data object, the MS2 spectrum, consists of a bag of peaks, where each peak is characterized by an *m/z* value and its associated intensity (Figure 1). The intensity is unitless but is monotonically related to the number of ions that generated the observed peak. The *m/z* value is measured with extremely high precision, often better than 10 parts-per-million (ppm), whereas the intensity is measured much less precisely. In practice, each spectrum contains around 100 peaks, with considerable variation (10–476 in the data analyzed here). A mass spectrometer produces MS2 spectra at a rate of *∼* 20–40 Hz, and a typical mass spectrometry run lasts 30–60 minutes, yielding on the order of 36,000–144,000 spectra in a single run.

The first analysis task that this data presents is the spectrum identification problem, in which each observed MS2 spectrum must be linked with the sequence of the peptide responsible for generating it. The primary constraint is that the peptide mass must lie within a specified tolerance of the observed precursor mass associated with the spectrum. This task is challenging because some of the expected b-ion and y-ion peaks may be missing, and some additional peaks may appear in the spectrum, created by losses of small molecular groups during fragmentation or by multiple cleavage events occurring on the same peptide. Spectra also contain noise, including experimental noise from the instrument as well as chemical noise produced by contaminants, other peptides, or non-peptide molecules.

In practice, spectrum identification is most commonly solved using database search, in which candidate peptides are selected from a given database of peptides [16]. The database typically contains all of the peptide sequences encoded in the genome of the species from which the sample is derived. *De novo* peptide sequencing, on the other hand, considers all possible peptide sequences and is useful for identifying sequences that arise from genetic variation, recombination in some immune system settings, vaccine design, antibody sequencing, or in the analysis of metaproteomic samples from many different types of microorganisms.

## 3 Related Work

Early *de novo* methods used heuristic search (Lutefisk [5]) or dynamic programming (PEAKS [6] and SHERENGA [7]) to score peptide sequences against each observed spectrum. The PepNovo algorithm [8] uses a similar dynamic programming approach but employs a probabilistic score function that takes into account various chemical and physical rules governing peptide fragmentation. This model is closely related to the hidden Markov model that is, to our knowledge, the first application of machine learning to the *de novo* peptide sequencing task [17]. A decade later, the Novor algorithm [9] achieved improved performance by using a decision tree as the score function in a dynamic programming algorithm.

The first deep neural network algorithm for *de novo* peptide sequencing, DeepNovo [2], combines two different network architectures—a convolutional neural network and a long short term memory (LSTM) network—each of which aims to predict the subsequent amino acid, given a spectrum and a peptide prefix. These two scores are combined in a dynamic programming procedure to yield the predicted peptide sequence. The recently described SMSNet algorithm [12] uses a network architecture similar to that of DeepNovo but also offers a post-processing step in which low-confidence amino acids are replaced by making use of a user-supplied peptide database. A competing method, pNovo 3 [11], works in three steps: (1) a traditional dynamic programming approach generates a set of candidate peptides for a given spectrum, (2) a previously described deep learning model, pDeep [18], predicts a theoretical spectrum for each candidate, and (3) a ranking support vector machine ranks the candidate peptides, based on features extracted by comparing the observed and theoretical spectra. Finally, PointNovo [10] is an improved version of DeepNovo which focuses specifically on handling high-resolution mass spectrometry data by using an order-invariant network architecture [19].

## 4 Methods

Transformers are highly capable of learning contextualized representations and modeling sequential data [20], with a variety of successful applications to biological sequences [21, 22]. In this context, *de novo* peptide sequencing can be formulated as a sequence-to-sequence learning problem where variable-length sequences of observed spectra peaks are translated into variable-length sequences of amino acids. The main contribution of this paper is to propose a transformer-based *de novo* peptide sequencing framework, Casanovo, which provides a unified solution to *de novo* peptide sequencing sub-tasks such as learning latent representations for spectra, spectrum processing and peptide sequence processing, which existing methods tackle separately through more complex modeling schemes.

### 4.1 Casanovo

Casanovo consists of a transformer encoder and decoder stack as described in [20], which are respectively responsible for learning latent representations of the input spectrum peaks and decoding the amino acid sequence of the spectrum’s generating peptide (Figure 2). The encoder takes *d*-dimensional spectrum peak embeddings as input and outputs *d*-dimensional latent representation vectors for each peak. Subsequently, the decoder takes as input these representations of prefix amino acids, coupled with a *d*-dimensional precursor embedding encapsulating precursor *m/z* and charge information, to predict the next amino acid in the peptide sequence. We discuss different aspects of our modeling strategy in detail below.

**Figure 2:**
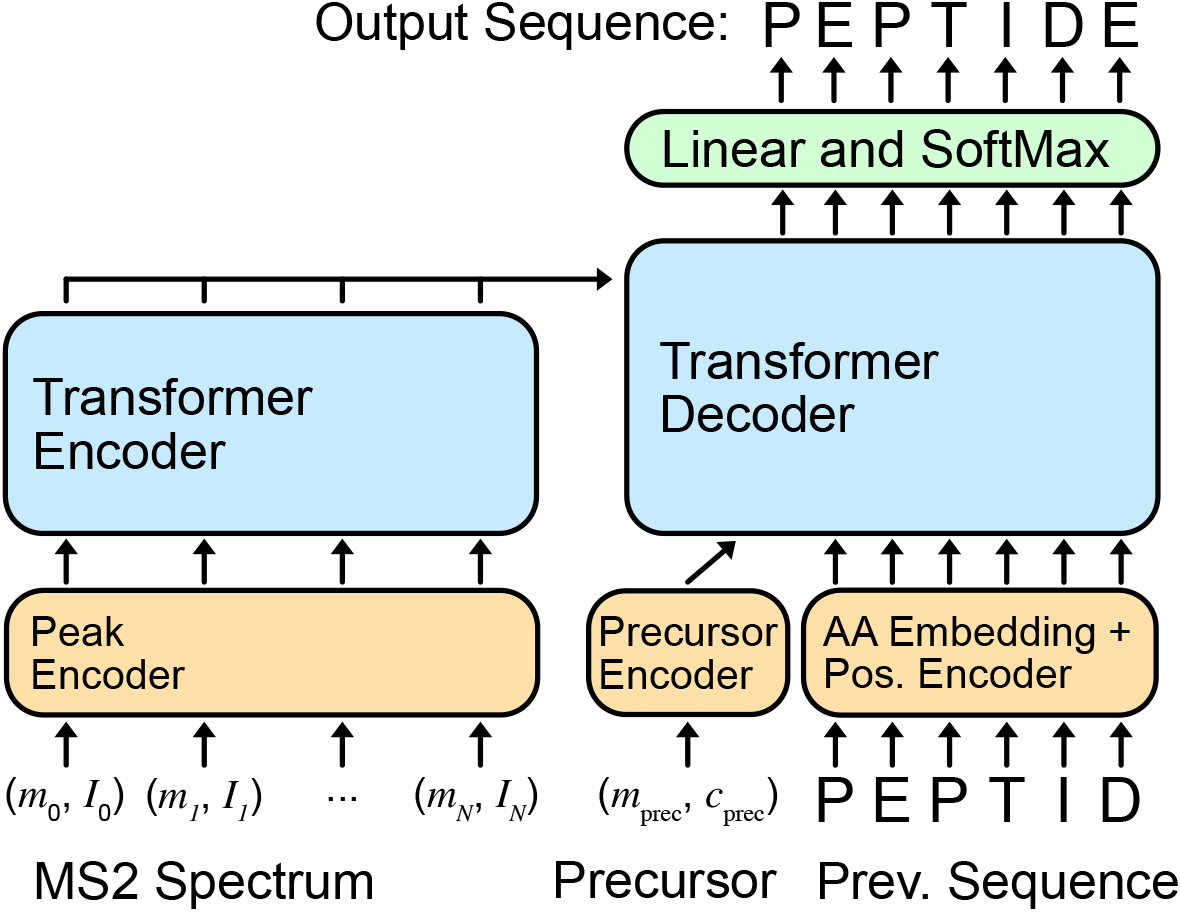
Casanovo model architecture with inputs and outputs.

#### 4.1.1 Input embeddings

Each spectrum 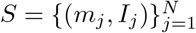 is a bag of peaks, where each peak (*m*_*j*_, *I*_*j*_) is a 2-tuple representing the *m/z* value and intensity of the peak. The *m/z* value and intensity are embedded separately before being summed to yield the input peak embedding. We use a fixed, sinusoidal embedding [20] to project each *m/z* value to a *d*-dimensional vector, the *m/z* embedding *f*. Specifically, we create the *m/z* embedding from an equal number of sine and cosine waveforms spanning the wavelengths from 0.001 to 10,000 *m/z*, where each feature in the embedding *f*_*i*_ is

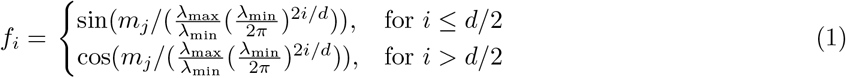

where *λ*_max_ = 10, 000 and *λ*_min_ = 0.001. These input embeddings provide a granular representation of high-precision *m/z* information and, similar to relative positions in the original transformer model [20], may help the model attend to *m/z* differences between peaks, which are critical for identification of amino acids in the peptide sequence. The intensity, which is measured with lower precision than the *m/z* value, is embedded by projection to *d* dimensions through a linear layer, after which the *m/z* and intensity embeddings are summed to produce the input peak embedding. We also experiment in Section 5.3 with encoding intensity using a fixed, sinusoidal position embedding and concatenating it with the *m/z* embedding.

Precursor information, used as input to the decoder, consists of the total mass *m*_prec_ *∈ R* and charge state *c*_prec_ *∈ {*1, …, 10*}* associated with the spectrum. We use the same sinusoidal position embedding as peak *m/z* ‘s for *m*_prec_; *c*_prec_ is embedded using an embedding layer, and the embeddings are summed to obtain the input precursor embedding. Preceding amino acids in the peptide sequence, another decoder input, are also encoded as the sum of an amino acid embedding and a sinusoidal position embedding of their position in the sequence.

#### 4.1.2 Training and inference strategy

Taking the previously described embeddings as input, the transformer outputs scores which are treated as a probability distribution over the amino acid vocabulary for the next position in the sequence at each decoding step. The amino acid vocabulary includes 20 canonical amino acids, post-translationally modified versions of three of them (oxidation of methionine and deamidation of asparagine or glutamine), plus a special stop token to signal the end of decoding, yielding a total of 24 tokens. During training, the decoder is fed the amino acid prefix for the ground truth peptide following the teacher forcing paradigm [23]. Cross-entropy between the model output probabilities and a binary matrix representing amino acid sequence of the ground truth peptide is minimized as the objective function. During inference, the highest scoring amino acid is predicted for each position in the sequence, and the decoder is fed its previous amino acid predictions at each decoding step. The decoding is finished either when the stop token is predicted or the pre-defined maximum peptide length of *l* = 100 amino acids is reached.

#### 4.1.3 Model and training hyperparameters

We train models with nine layers, embedding size *d* = 512, and eight attention heads, yielding a total of *∼* 47M model parameters. A batch size of 32 spectra and 10^*−*5^ weight decay is used during training, with a peak learning rate of 5 *×* 10^*−*4^. The learning rate is linearly increased from zero to its peak value in 100k warm-up steps, followed by a cosine shaped decay. Models are trained on 2 RTX 2080 GPUs for 30 epochs, which takes approximately two days, and model weights from the epoch with the lowest validation loss were selected for testing. These model hyperparameters—number of layers, embedding size, number of attention heads, and learning rate schedule—are used for all downstream experiments unless otherwise specified.

#### 4.1.4 Precursor *m/z* filtering

A critical constraint in *de novo* peptide sequencing requires the relative difference between total mass of the predicted peptide *m*_pred_ and the observed precursor mass *m*_prec_ of the spectrum to be less than a threshold value *ϵ* (specified in ppm) for the predicted sequence to be plausible: 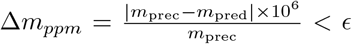 In addition to providing precursor information as an input for the model to learn from, we filter out peptide predictions that do not satisfy the above constraint. The threshold value *ϵ* is a property of the mass spectrometer that the data is collected with, and hence is known at inference time. Accordingly, we choose *ϵ* based on the precursor mass error tolerance used in the database search to obtain ground truth peptide sequences for the test data.

Casanovo’s source code and trained model weights are available as open-source under the Apache 2.0 license at https://github.com/Noble-Lab/casanovo.

### 4.2 Data set

To evaluate the performance of Casanovo and compare it with state-of-the-art *de novo* peptide sequencing methods, we use the nine-species benchmark data set and evaluation framework first introduced by [2] and used in several subsequent studies [12, 10]. This data set combines a total of about 1.5 million mass spectra from nine different experiments, each using the same instrument to analyze peptides from a different species. Based on database search identification using the standard false discovery rate (FDR) of 1%, each spectrum comes with an assigned peptide sequence which is treated as ground truth to train and evaluate the methods. With approximately 300,000 unique peptide sequences in the data set, each sequence has around five spectrum instances on average, but around 40% of all peptide sequences have a single spectrum associated with them. Following [2], we employ a leave-one-out cross validation framework where we train a model on eight species and test on the held-out species for each of the nine species in the data set. In each case, we split the training set 90/10 for training and validation. This cross-species evaluation framework allows for testing the model on never-before-seen peptide samples, because the peptides in the training set are almost completely disjoint from the peptides of the held-out species. To illustrate this point, among the *∼* 26,000 unique peptide labels associated with the human spectra in the test data, only 7% overlap with the *∼* 250,000 unique peptide labels associated with spectra from the other eight species. Cross-species testing is particularly important for *de novo* sequencing models because most practical applications of *de novo* sequencing require models to perform well on spectra with never-before-seen peptide sequences.

### 4.3 Evaluation metrics

We use precision calculated at the amino acid and peptide levels [6, 8, 2] as a function of coverage over the test set as performance measures to evaluate the quality of a given model’s predictions. In each case, for each spectrum we compare the predicted sequence to the ground truth peptide from the database search. Following [2], for the amino acid-level measures we first calculate the number 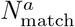 of matched amino acid predictions, defined as all predicted amino acids which (1) differ by *<*0.1 Da in mass from the corresponding ground truth amino acid, and (2) have either a prefix or suffix that differs by no more than 0.5 Da in mass from the corresponding amino acid sequence in the ground truth peptide. We then define amino acid-level precision 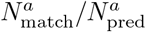, where 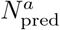 is the number of predicted amino acids. For peptide predictions, a predicted peptide is considered a correct match if all of its amino acids are matched. Among a collection of 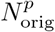 spectra, if our model makes predictions on a subset of 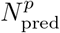 and correctly predicts 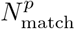 peptides, we define coverage as 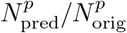 and peptide-level precision as 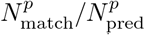. To plot a precision-coverage curve, we sort predictions by the confidence score provided by the model. Amino acid-level confidence scores are obtained by applying a softmax to the output of the transformer decoder, which is a proxy for the probability of each predicted amino acid to occur in the given position along the peptide sequence. Casanovo directly outputs amino acid-level confidence scores, and we use the mean score over all amino acids as a peptide-level confidence score.

## 5 Results

### 5.1 Casanovo outperforms state-of-the-art methods

We begin by using the previously described experimental setup and evaluation metrics (Sections 4.2–4.3) to evaluate Casanovo’s performance relative to two state-of-the-art neural network-based methods, DeepNovo [2] and PointNovo [10]. In this comparison, peptide-level performance measures are the primary quantifier of the sequencing model’s practical utility, since the goal is to assign a complete peptide sequence to each observed spectrum. To characterize the performance of DeepNovo and PointNovo, we rely on the pre-trained weights of the former and the published results of the latter [10], since neither PointNovo’s pre-trained weights nor its predictions for the benchmark data set are available.

At the peptide level, Casanovo substantially outperforms both previous methods across all species, with a mean improvement of 0.373 and 0.310 in precision relative to DeepNovo and PointNovo, respectively, at a mean coverage of 0.60 (Table 2). Even when precursor *m/z* filtering is turned off and the model is forced to make a prediction for all spectra, i.e. coverage is 1.0, Casanovo shows a mean improvement of 0.076 and 0.013 relative to DeepNovo and PointNovo, respectively. Indeed, the peptide-level precision-coverage curves (Figure 3) show that Casanovo consistently outperforms DeepNovo over a range of peptide confidence thresholds. This trend is also reflected by the area under the curve (AUC) metric (Figure 3), with Casanovo outperforming DeepNovo by 0.131 on average.

**Table 2:**
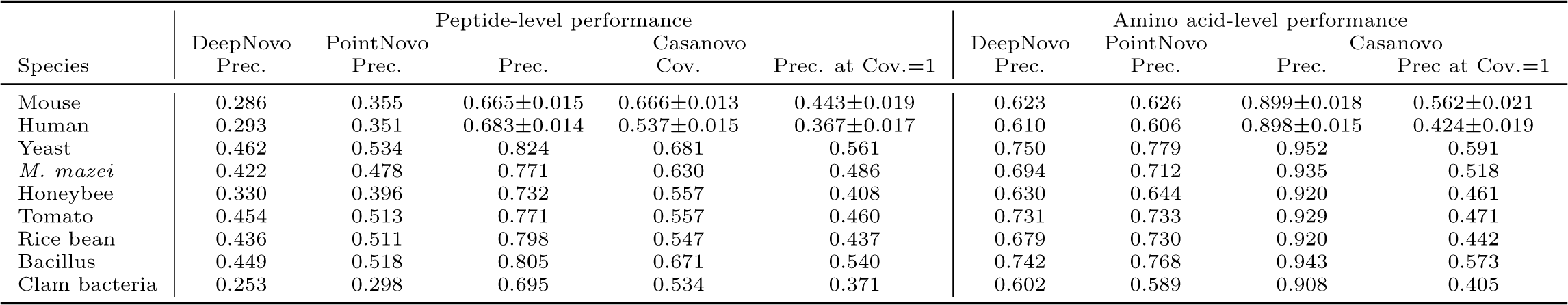
Empirical comparison of Casanovo, DeepNovo and PointNovo. The table lists the peptide-level and amino acid-level precision of three competing models and coverage of Casanovo with precursor *m/z* filtering on all nine benchmark cross-validation folds. Each fold’s test set contains spectra from a single species, with nearly disjoint sets of peptides between species. For cross-validation folds corresponding to mouse and human, five models were trained with different random initializations. For these species, we report standard deviation of the performance measures.

**Figure 3:**
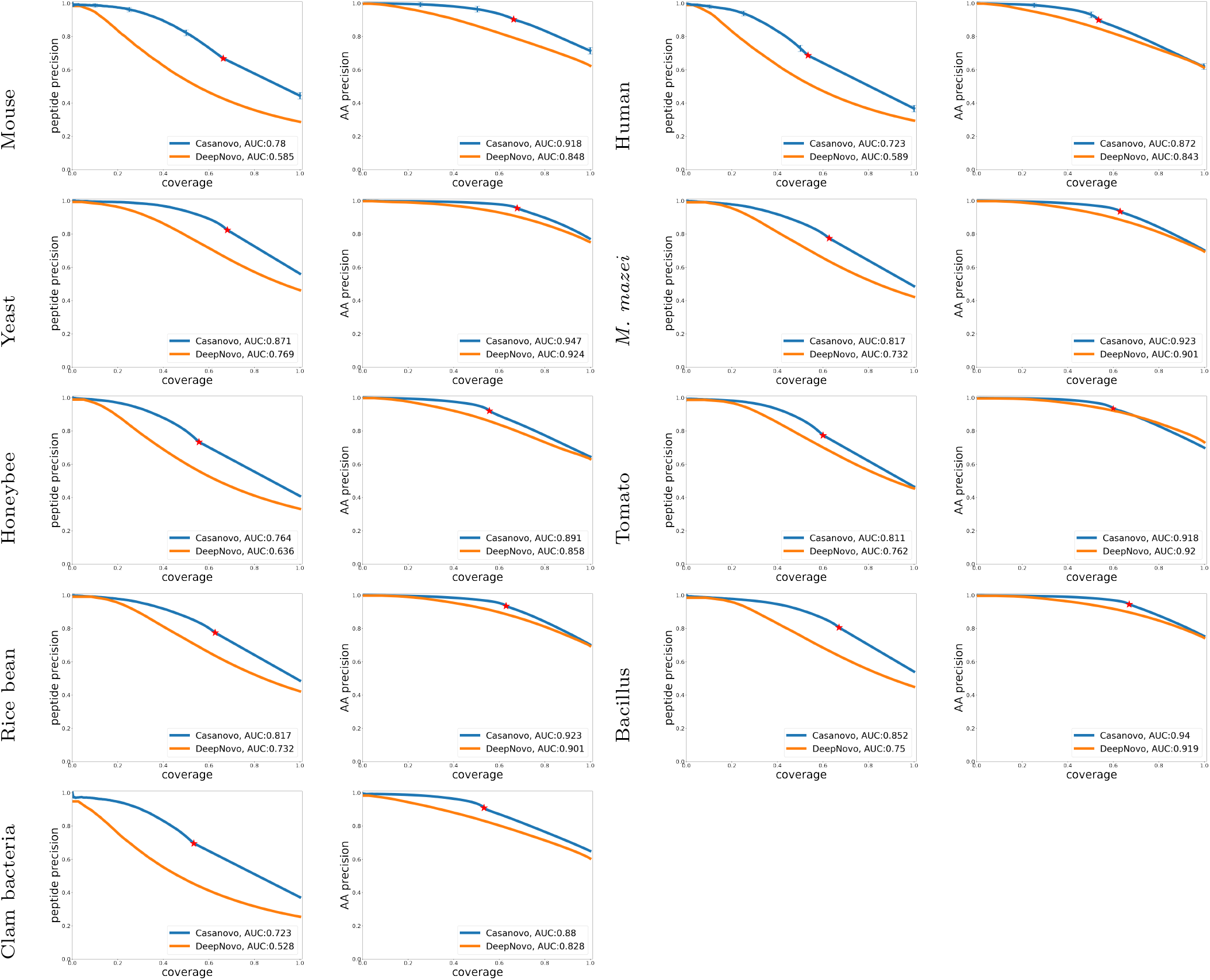
Precision-coverage curves for Casanovo and DeepNovo. Curves are shown for each species (two per row) at the peptide level (left sub-column) and amino acid level (right sub-column). Curves are computed by sorting predicted peptides according to their confidence scores. For the amino acid level curves, all amino acids within a given peptide receive equal scores. For Casanovo at both amino acid and peptide level, all peptides that pass the precursor *m/z* filtering are ranked above peptides that do not pass the filter, and similarly for all amino acids from peptides that pass the precursor *m/z* filtering versus those that do not pass the filter. The boundary between unfiltered and filtered entries is indicated by a red star on each curve. Error bars are provided for human and mouse species, for which five models were trained with different random initializations.

Similarly, at the amino acid level, Casanovo outperforms DeepNovo and PointNovo, particularly in the high-precision portions of the curves. As expected, the precursor *m/z* filtering, which prioritizes predicting full peptide sequences with high precision over partially correct peptide predictions, yields better overall precision at the cost of reduced precision at full coverage. In all nine species, the point on the Casanovo curve corresponding to the filter lies above the DeepNovo precision-coverage curve, and in eight of the nine species

Casanovo’s AUC exceeds DeepNovo’s. We further discuss the effects of precursor *m/z* filtering in Section 5.2. Complementing its improved *de novo* peptide sequencing performance, Casanovo achieves these results with fewer model parameters (47 M) than DeepNovo (86 M). (The number of parameters and model dimensions were not reported for PointNovo.) Casanovo also runs inference at a faster rate of 119 spectra/s on an RTX 2080 compared to DeepNovo’s 36 spectra/s and PointNovo’s reported 20 spectra/s on an RTX 2080 Ti (a comparatively faster GPU) [10].

### 5.2 Precursor *m/z* post-processing

One of the key components of DeepNovo, and its successor PointNovo, is a post-processor that uses the knapsack dynamic programming algorithm to ensure that the mass of the predicted peptide is close to the observed precursor mass. Ablation experiments in the DeepNovo experiment paper show that removing this component leads to a decrease in peptide-level precision of 12.4% (averaged over test sets) [2]. Accordingly, we tested three variants of Casanovo on the yeast species test set: no post-processor, the knapsack post-processor, and our simple *m/z* filter (Figure 4).

**Figure 4:**
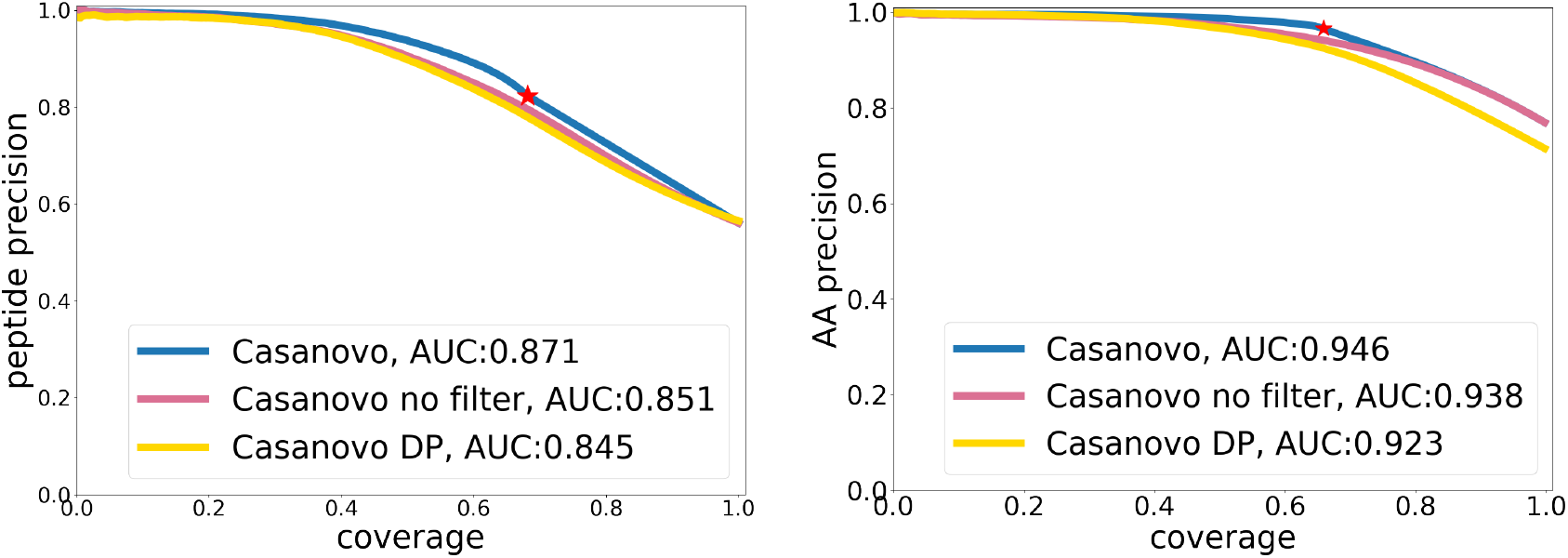
Precision-coverage curves for Casanovo models with different post-processors. Standard Casanovo model with simple filter outperforms both no filter and dynamic programming post-processor on the yeast test set, where we see that the effect of the filter is to boost precision along the entire curve.

At the peptide level, we observe a much smaller benefit from the knapsack algorithm—an increase in the peptide-level precision from 0.561 to 0.565 when models are compared at full coverage—than was reported in the DeepNovo paper. On the other hand, a comparison of precision-coverage curves indicates that the knapsack algorithm hurts the AUC metric and precision at most coverage values.

At the amino acid level, the effects of these two post-processors is different. Relative to Casanovo with no post-processing, applying the knapsack algorithm yields a small decrease in precision at full coverage (0.769 *→*0.726), whereas adding the precursor *m/z* filtering yields much higher precision (0.769 *→*0.965) which decreases substantially when the coverage is extended to all spectra (0.769 *→*0.576). Similar to the peptide level, the standard Casanovo model with the simple filter consistently yields the highest precision for differenct values of coverage as well as the largest AUC.

To better understand these results, we performed a qualitative review of the predictions from the three models. This analysis suggests that incorrect amino acid predictions in earlier decoding steps cause the post-processor to discard correct amino acids from among options in later decoding steps, leading to a drop in amino acid-level performance. This observation is supported by the plot of the precision-coverage curve with and without the precursor *m/z* filter (Figure 4), where we see that the effect of the filter is to boost precision along the entire curve.

### 5.3 Peak embeddings and loss function

Finally, we train three additional variants of Casanovo, none of which provides any performance improvements over the standard model (Table 3). The first variant uses a focal loss function, adopted from [10], instead of cross entropy. We also investigated two alternate methods of peak embedding. The first, also adopted from PointNovo, replaces summation of *m/z* and *I* embeddings with direct multiplication of the *I* value and the *m/z* embedding. The second peak embedding strategy implements a sinusoidal encoding for *I*, similar to the *m/z* embedding, although using only 32 dimensions, and concatenates *I* with the *m/z* embeddings instead of summing the two.

**Table 3:**
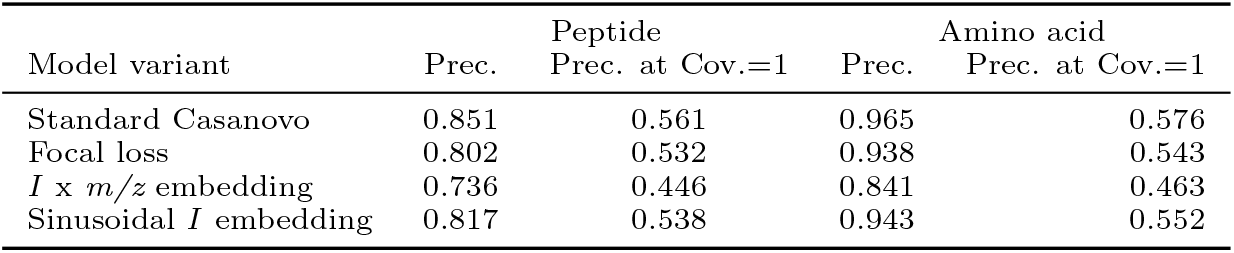
Performance comparison of different Casanovo variants. All results are for the yeast test set.

## 6 Discussion

Prior work in *de novo* peptide sequencing has used deep learning models that combine separate neural network architectures followed by complex post-processing steps. Our approach, Casanovo, leverages the transformer architecture to produce a unified solution to translate mass spectra directly into peptide sequences, without resorting to discretization of the spectrum *m/z* axis and without complex post-processing. We find that Casanovo achieves state-of-the-art performance on the standard benchmark data set, with fewer model parameters compared to existing methods.

Casanovo’s inference speed is fast enough to allow real time *de novo* sequencing, i.e., sequencing at the speed that the mass spectrometer generates spectra, raising the possibility of helping guide mass spectrometry experiments as they are being conducted [9]. In practice, real-time search results can be useful for making decisions about peptide elution order [24], improving the accuracy of stable isotope labeling [25], post-translational modification site localization [25], or deciding whether to trigger an MS3 (secondary fragmentation) scan [26].

Casanovo improves substantially over the previous state of the art in terms of peptide-level precision, but this leaves a significant portion of the test spectra without plausible predictions. Clearly, exploring methods to find good predictions for these spectra is an avenue for future research. To explore the potential benefit of such an approach, we combined Casanovo and DeepNovo predictions by inserting DeepNovo predictions whenever the *m/z* filter eliminates a Casanovo prediction. The resulting model achieves up to 10% higher peptide precision than Casanovo and exceeds the previous state-of-the-art method, PointNovo, on all evaluation metrics across species. This observation suggests that precursor information should be included as a stronger prior in modeling mass spectra. A straightforward approach might involve choosing among a larger set of peptide candidates generated by beam search during inference, with a constraint on the predicted mass. Alternatively, Casanovo’s loss function could be modified to penalize peptide predictions which do not match the precursor mass.

## Acknowledgement

This work is in part supported by CCF-2019844 as a part of the NSF Institute for Foundations of Machine Learning (IFML).

